# Cryo-EM reveals a right-handed double-helix dimer architecture of PCDH15 critical for mechanotransduction

**DOI:** 10.64898/2026.03.02.709101

**Authors:** Xiaoping Liang, Roshan Pathak, Xufeng Qiu, Lucas Dillard, Edward C. Twomey, Ulrich Müller

## Abstract

Tip links connect the stereocilia of mechanosensory hair cells in the inner ear and transmit force onto mechanotransduction (MET) channels. Tip links consist of protocadherin 15 (PCDH15) and cadherin 23 (CDH23), which assemble into an extracellular filament approximately 150 nm in length. Rare freeze-etched electron microscopy (EM) images have suggested that tip links could be right-handed double helices in vivo, but direct structural evidence has been lacking. Using cryo-EM we determined the structure of a large part of the extracellular PCDH15 domain. Two PCDH15 molecules form a parallel cis dimer stabilized by several dimerization interfaces, including two strand crossovers and two parallel contacts, yielding a right-handed double helix. Functional studies show that mutations in PCDH15 dimerization-domains impair MET. Our results establish the molecular foundation for how PCDH15 forms a right-handed double helix to enable mechanical sensing.

## Introduction

Hair cells of the vertebrate inner ear are specialized mechanosensory cells that transduce mechanical stimuli arising from sound waves and head movement into electrical signals (Fig. 1A). Hair cells contain at the apical surface a bundle of stereocilia that are organized in rows of decreasing height. Deflection toward the tallest row of stereocilia opens MET channels positioned at the base of tip links, the fine extracellular filaments aligned with the bundle’s axis of sensitivity, thereby coupling mechanical force to channel gating (1, 2).

**Figure 1.**
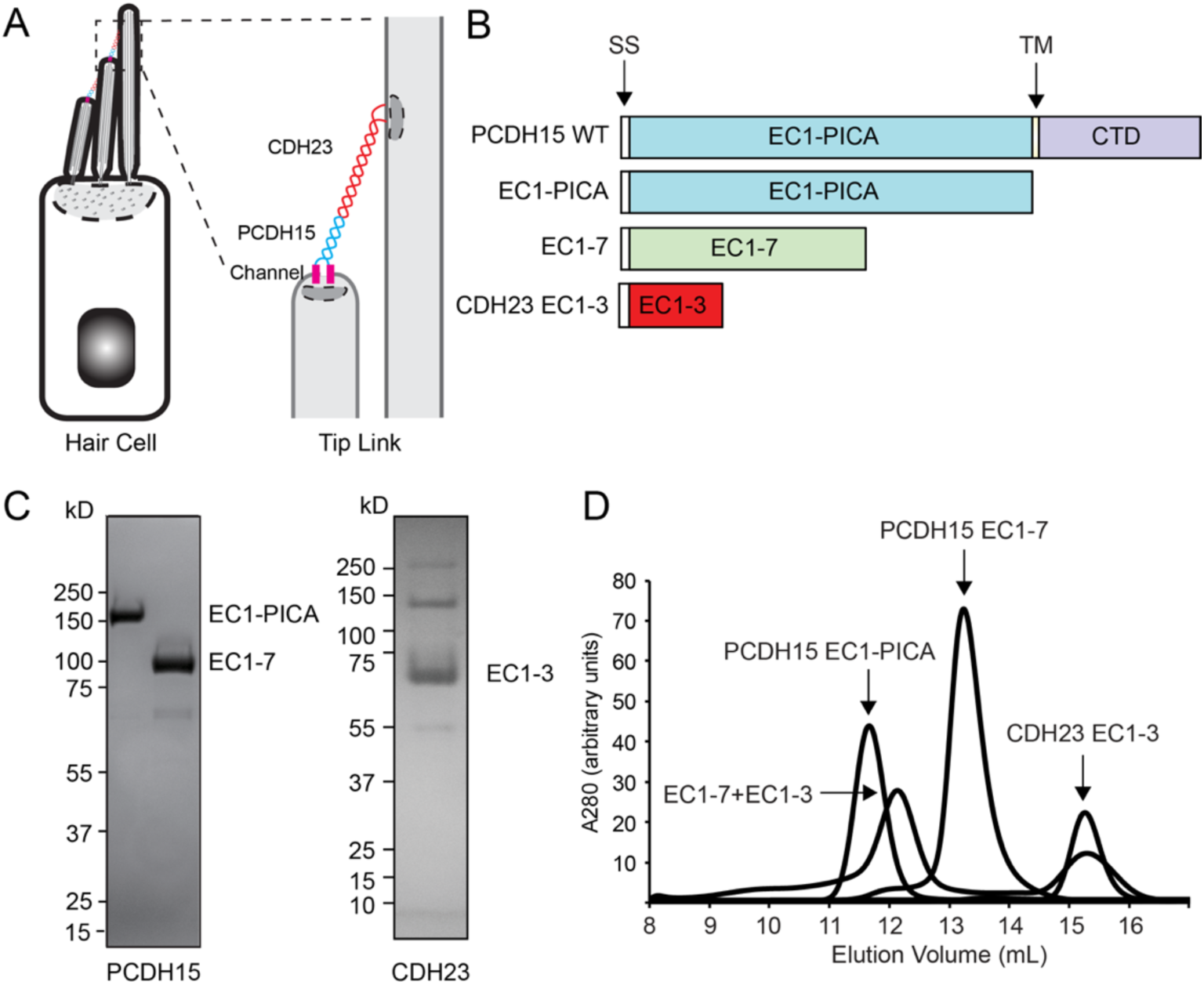
Purification of the recombinant extracellular PCDH15 domain. (A) Diagram of a hair cell highlighting the tip link complex consisting of homodimers of PCDH15 (blue) and CDH23 (red) at the tips of stereocilia. (B) Diagram of PCDH15 and CDH23 expression constructs (SS, signal sequence; TM, transmembrane domain; CTD, C-terminal cytoplasmic domain). (C) Purified extracellular domain fragments were separated on SDS-PAGE gels and visualized by Coomassie staining. (D) Size exclusion chromatograms of the indicated purified recombinant proteins monitored by absorbance at 280 nm. PCDH15-EC1-PICA, PCDH15-EC1-7, and CDH23-EC1-3 each elute as single, symmetric peaks. Upon mixing, PCDH15-EC1-7 and CDH23-EC1-3 co-elute at an earlier elution volume consistent with complex formation.

Biochemical evidence indicates that tip links are heteromeric filaments formed by PCDH15 and CDH23, measuring approximately 150 nm in length (3). Unlike classical cadherins, which typically have five extracellular cadherin (EC) domains, the longest splice variants of PCDH15 and CDH23 have 11 and 27 EC domains, respectively (4–8). Crystal structures show that individual EC domains of tip-link cadherins adopt classical cadherin folds (9–18). However, unlike those of classical cadherins, their inter-EC linkers do not uniformly bind three Ca^2+^ ions (9, 16, 18), implying segments of reduced rigidity that may be functionally important for MET. Furthermore, PCDH15 and CDH23 form stable cis homodimers that engage in trans to form the lower and upper halves of the tip link (3), an adhesion mode distinct from classical cadherins, which rely on weaker cis interactions and primarily homophilic trans adhesion via EC1 (19–21). In contrast, tip-link cadherins engage in more extensive heterophilic EC1-EC2 trans interactions (10, 13, 18) likely conferring enhanced mechanical stability to the tip-link trans-bond. Near the membrane, PCDH15 and CDH23 contain non-classical domains, variously termed MAD, EL, SEA-like, or PICA (11, 12, 22, 23), further distinguishing their architecture from classical cadherins and likely reflecting adaptations for MET.

Freeze-etch electron microscopy (EM) images suggest that tip links may adopt a right-handed double helix (24), but direct evidence has been limited. While structural studies resolved details of several PCDH15/CDH23 EC domains and of the PCDH15 PICA domain, including dimerization motifs at EC2-EC3 and within the PICA domain (9–18, 23), short fragments could not reveal the global helical organization of tip links. The elongated shape of full-length PCDH15 and CDH23 also complicates cryo-EM analysis.

Here, we resolved the structure of PCDH15 EC1-EC7 domain by cryo-EM, revealing a right-handed double helix formed by two PCDH15 molecules. Dimerization is mediated by interfaces at EC2-EC3, EC4-EC5, and EC6-EC7. The EC2-EC3 interface matches prior reports (10, 12) while EC4-EC5 and EC6-EC7 are newly defined here. Strand crossovers occur at EC2-EC3 and EC6-EC7, with parallel alignment at EC4-EC5 and in the PICA region (23). We also show that PCDH15 mutations within dimer interfaces reduce MET in hair cells.

## Results

### Analysis of the structure of the extracellular PCDH15 domain using cryo-EM

To determine the atomic structure of the extracellular PCDH15 domain, we engineered PCDH15 constructs lacking the transmembrane and cytoplasmic domains (Fig. 1B). We also inserted C-terminal FLAG or streptavidin tags and expressed the constructs in HEK293 cells to ensure proper folding and glycosylation. We purified full-length extracellular PCDH15 (PCDH15-EC1-PICA; aa 1-1381) and a truncated EC1-EC7 construct (PCDH15-EC1-7; aa 1-824) by affinity chromatography using streptavidin followed by size-exclusion chromatography. On SDS-PAGE gels, both proteins resolved as a distinct band of the predicted size (Fig. 1C). Both proteins were stable each resulting in a monodispersed peak when analyzed by size exclusion chromatography (SEC) (Fig. 1D). When PCDH15-EC1-7 was mixed with a fragment of CDH23 consisting of the first three EC domains (and containing C-terminal Fc and streptavidin tags; CDH23-EC1-3) (Fig. 1B), which contain the PCDH15 binding site (10, 18), the two fragments co-eluted at an earlier elution volume from the SEC column (Fig. 1D), consistent with binding to each other and suggesting that the CDH23 binding domain was properly folded in PCDH15-EC1-7.

We determined the structure of the extracellular PCDH15 domain by cryo-EM (Supplementary Fig. 1; Supplementary Table 1). PCDH15-EC1-PICA appeared dimeric by 2D class averages (Fig. 2A,C) but proved conformationally heterogeneous, limiting resolution. In contrast, PCDH15-EC1-7 yielded well-distributed dimeric particle (Fig. 2B,C). The elongated PCDH15-EC1-PICA and PCDH15-EC1-7 particles displayed a distinct crossed arrangement with a bifurcated “tail”-like structure (Fig. 2A-C). PCDH15-EC1-PICA also had a globular density (Fig. 2A, box), which was not present in PCDH15-EC1-7 (Fig. 2B), likely formed by the PICA domains of the two strands. We determined the structure of PCDH15-EC1-7 at an overall resolution of approximately 3.8 Å, with local refinements of 3.43 Å (EC1-EC2), 3.40 Å (EC3-EC5), and 3.83 Å (EC6-EC7) (Supplementary Fig. 2). The dimer spanned approximately 320 Å, formed a right-handed double helical turn, and had a ∼120 Å diameter, consistent with EC-domain dimensions. One end of the homodimer formed a scissor-like crossing and separation of the two strands, consistent with previous crystallographic studies that have shown a similar strand separation at EC1 and EC2 of PCDH15 (10, 12, 18). The remaining EC repeats adopted a twisted, braided configuration along the longitudinal axis.

**Figure 2.**
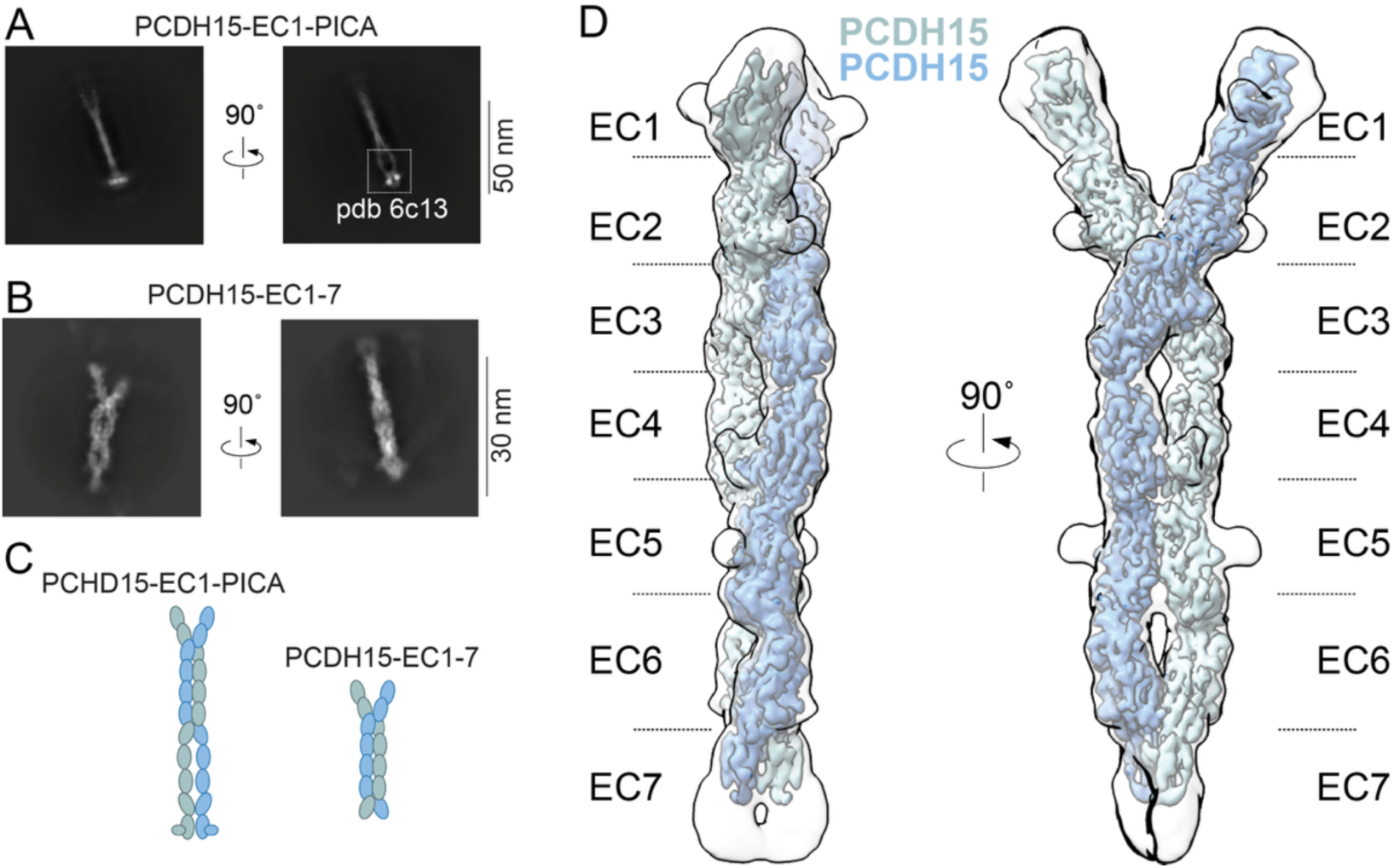
Cryo-EM structure of PCDH15-EC1-7. (A,B) Representative cryo-EM micrographs of recombinant PCDH15-EC1-PICA (A) and PCDH15-EC1-7 (B). The right panel shows the same particle view as on the left but rotated by 90°. Scale bar: A, 50 nm, B, 30 nm. (C) Diagram of the dimeric recombinant molecules, with ovals representing EC domains. (D) Cryo-EM structure of the PCDH15-EC1-7 homodimer. Protomers are colored separately (light blue and teal), and individual EC domains (EC1–EC7) are indicated.

### Domain fitting and analysis of the PCDH15 dimer interface

We fitted the map of EC1-7 to previously published X-ray structures of PCDH15 fragments (9–13). EC1-EC6 fitted well into the map with a partial fit for EC7 (Fig. 3). We identified extensive lateral contacts between PCDH15 strands at EC2-EC3, consistent with our previous X-ray crystallographic data (12), and at EC4-EC5 and EC6-EC7. EC2-EC3 and EC6-EC7 formed strand crossovers. EC4-EC5 domains aligned nearly parallel, with contributions from EC4 loop regions and EC5 β-sheets to lateral interactions (Fig. 3). Prior work also indicates a lateral interaction between PICA domains (23), adding a membrane-proximal interface.

**Figure 3.**
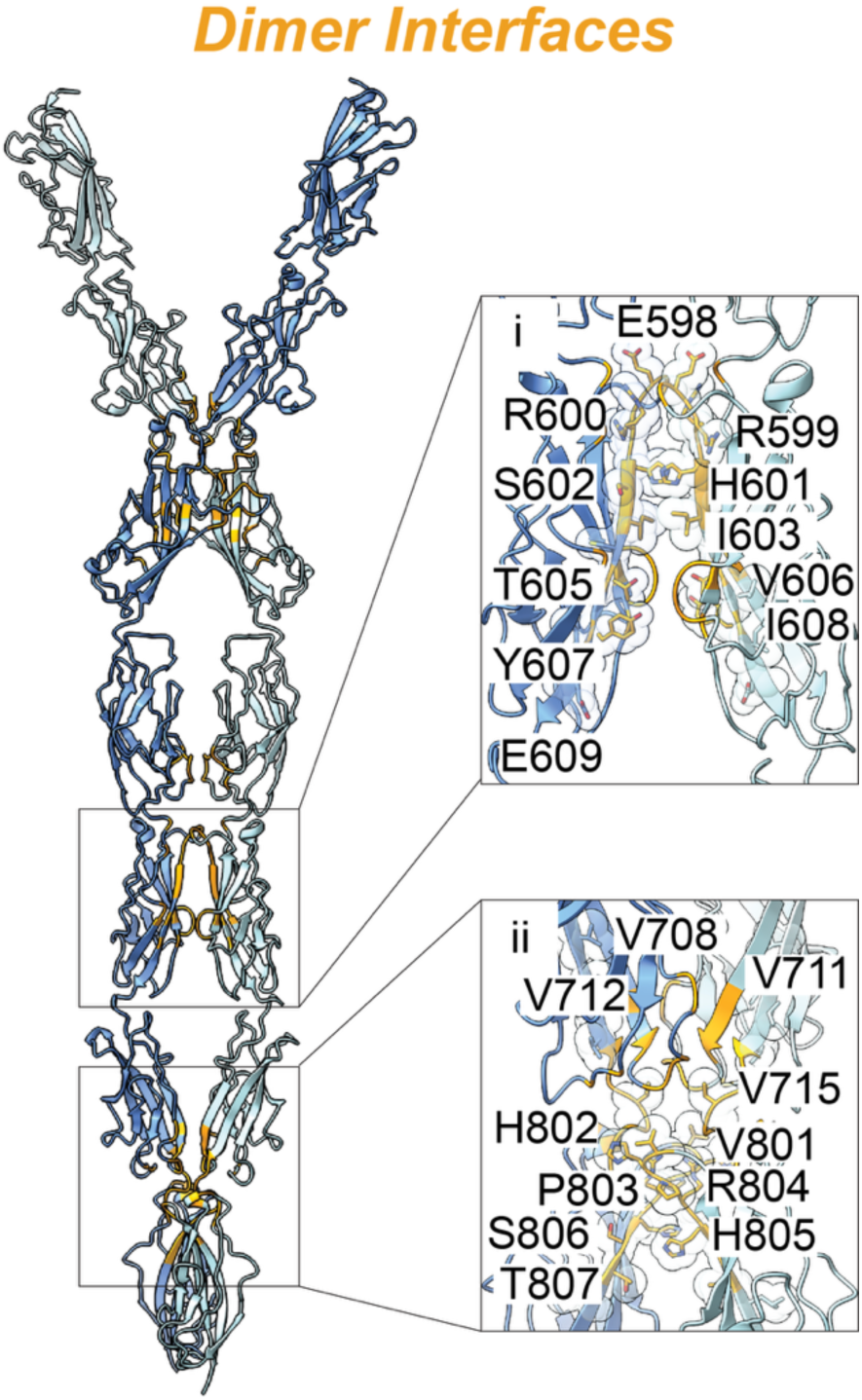
PCDH15 dimer interfaces. Structural details highlighting protomer-protomer interactions at EC4-EC5 and EC6-EC7. Interface residues are depicted in stick representation.

### Identification of mutations that disrupt PCDH15 dimerization

We previously confirmed a dimerization interface at the EC2-EC3 linker by mutational analysis and biochemical experiments (12). To validate the newly identified interfaces at EC4–EC5 and EC6-EC7 (Fig. 3), we designed mutant constructs for biochemical characterization (Fig. 4A). Guided by our structural data, we generated three PCDH15-EC1-PICA deletion constructs with C-terminal HA tags: Δ598-609 (affecting EC5), Δ710-720 (EC6), and Δ801-809 (EC7). Each HA-tagged construct was co-expressed in HEK293 cells with PCDH15-EC1-PICA bearing a C-terminal FLAG tag. We immunoprecipitated PCDH15-EC1-PICA-FLAG using anti-FLAG antibodies and probed for co-precipitated HA-tagged PCDH15-EC1-PICA by western blot (Fig. 4B). Densitometric quantification from at least three independent experiments showed that wild-type PCDH15-EC1-PICA-HA robustly co-immunoprecipitated with PCDH15-EC1-PICA-FLAG, whereas all three deletion mutants caused a significant reduction in co-immunoprecipitation efficiency (Fig. 4B). These results indicate that multiple interfaces contribute to the stabilization of the PCDH15 homodimer.

**Figure 4.**
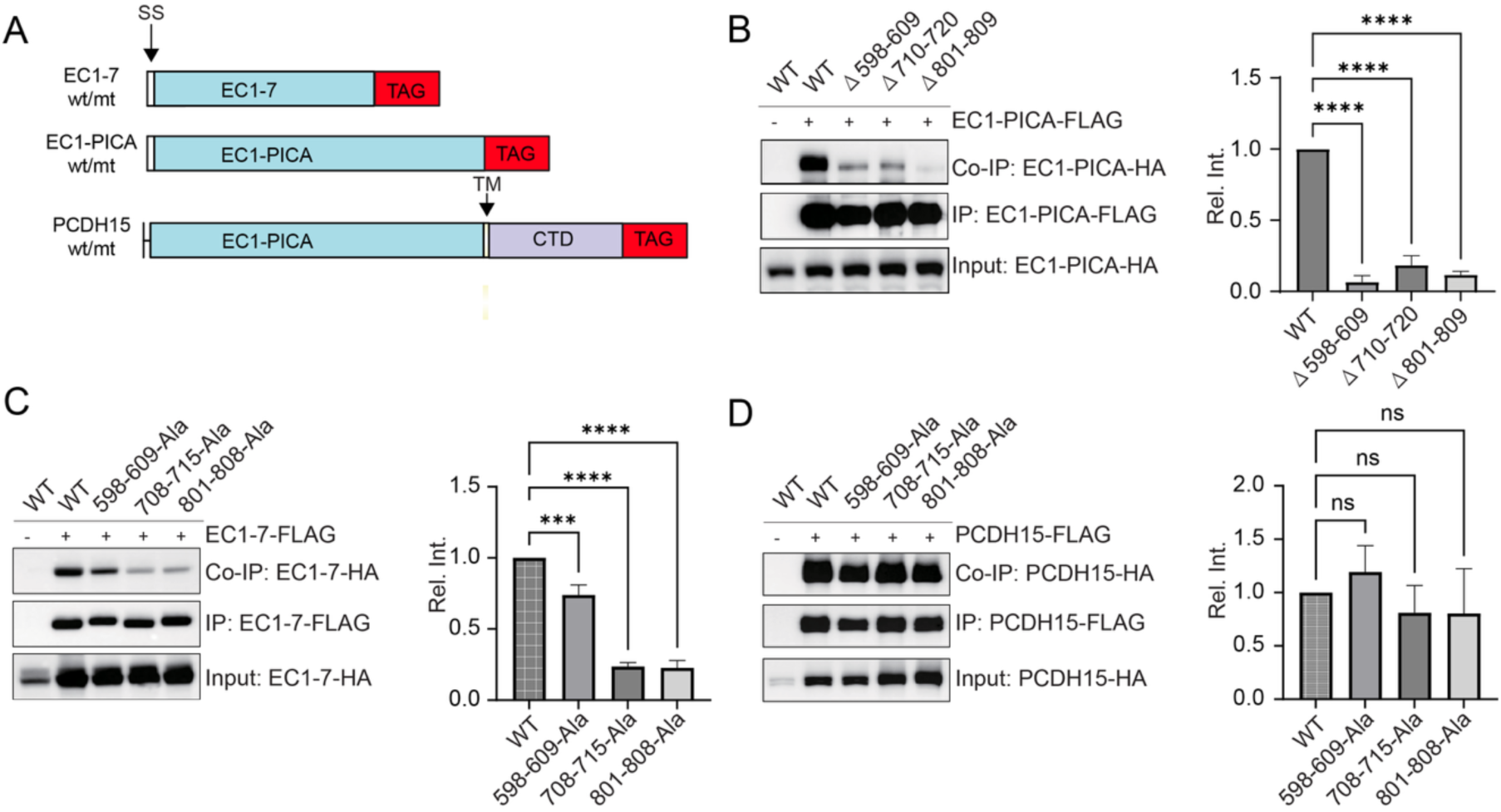
Analysis of EC4-EC5 and EC6-EC7 homodimer interfaces by mutagenesis and co-immunoprecipitations. (A) Schematic depicting constructs used for experiments (SS, signal sequence; TM, transmembrane domain). (B-D) HEK293 cells were transfected with the constructs indicated on top of each panel. Immunoprecipitations (IPs) were carried out with FLAG-conjugated beads, followed by Western blotting with antibodies to HA (upper rows, Co-Immunoprecipitation (CoIP); middle rows, Immunoprecipitation (IP); lower rows, input). Quantification of CoIP results from at least 3 independent experiments. (Rel. Int., Relative Intensity; mean ± SEM; Student’s t-test; n.s. p>0.05, *** p<0.001, **** p<0.0001).

Because deletions can disrupt the overall protein backbone and confound interpretation, we next generated point mutants that preserved the backbone but altered key residues on the predicted dimerization surfaces by substituting them with ALA. The point mutants were: EC5 surface (residues 598-609 → ALA), EC6 surface (708-715 → ALA; 708,710,711,712 and 715 were altered to ALA to disrupt lateral hydrophobic interactions; 709, 711 and 714 were unchanged to preserve Ca^2+^ binding residues), and EC7 surface (801-808 → ALA) (Fig. 4C). We avoided conserved Ca^2+^-binding residues in the EC linker regions. These alanine-substitution mutants significantly reduced dimerization of the PCDH15-EC1-7 fragment, as assessed by co-immunoprecipitation (Fig. 4C).

To assess the impact of these point mutations in the context of full-length PCDH15 including the transmembrane (TM) and cytoplasmic domains (CTD), we introduced the same substitutions into full-length PCDH15 constructs. For our experiments, we used the PCDH15-CD2 splice variant, which is present at tip links and promotes MET (12, 25, 26). Surprisingly, dimerization of full-length PCDH15 was not noticeably affected by any single mutant (Fig. 4D). Thus, when all four extracellular dimerization motifs as well as the TM and CTD domains of PCDH15 are present, disrupting a single interface is insufficient to abolish dimer formation, at least under the conditions of our co-immunoprecipitation assay. The findings also suggest that the TM and/or CTD domains contribute to stabilizing PCDH15 homodimers.

### Mutations that disrupt PCDH15 homodimerization impair MET in cochlear hair cells

We have previously shown that injectoporation is an effective procedure to deliver PCDH15 expression vectors into hair cells and to assess MET by monitoring fluorescence changes of the co-expressed Ca^2+^ sensor G-CaMP3 after hair-bundle stimulation with a fluid jet (Fig. 5A) (12, 27–29). Our prior work also established that mutations in the EC2-EC3 dimerization interface impair MET (12). To determine whether the EC4-EC5 and EC6-EC7 interfaces are also functionally important, we injectoporated full-length wild-type and ALA-substitution PCDH15-HA constructs together with G-CaMP3 into hair cells from PCDH15-deficient homozygous Ames-waltzer *av3J/av3J* mice (4) and measured MET responses. All three ALA-substitution constructs localized to stereocilia of OHCs, as shown by immunofluorescence with HA-specific antibodies, similar to wild-type PCDH15-HA (Fig. 5B). Injectoporated cells were identified by the GFP signal from co-expressed G-CaMP3. Hair bundles were stimulated with a fluid jet using three consecutive pulses of increasing duration (0.1, 0.3 and 0.5 s). As previously reported, hair cells from *av3J/av3J* mice showed no significant fluorescence increase after mechanical stimulation (12). Re-expression of wild-type PCDH15 restored a robust G-CaMP3 fluorescence response to bundle deflection (Fig. 5C). In contrast, OHCs expressing two of the three ALA-substitution mutants (residues 598-609 → ALA; 801-808 → ALA) exhibited significantly reduced fluorescence increases in response to mechanical stimulation, with the third mutation (708-715 → ALA) being highly variable but showing a trend towards reduced function (Fig. 5C). Together, these data indicate that weakening cis-homodimerization at the EC4-EC5 and EC6-EC7 interfaces compromises PCDH15 function in hair-cell MET.

**Figure 5.**
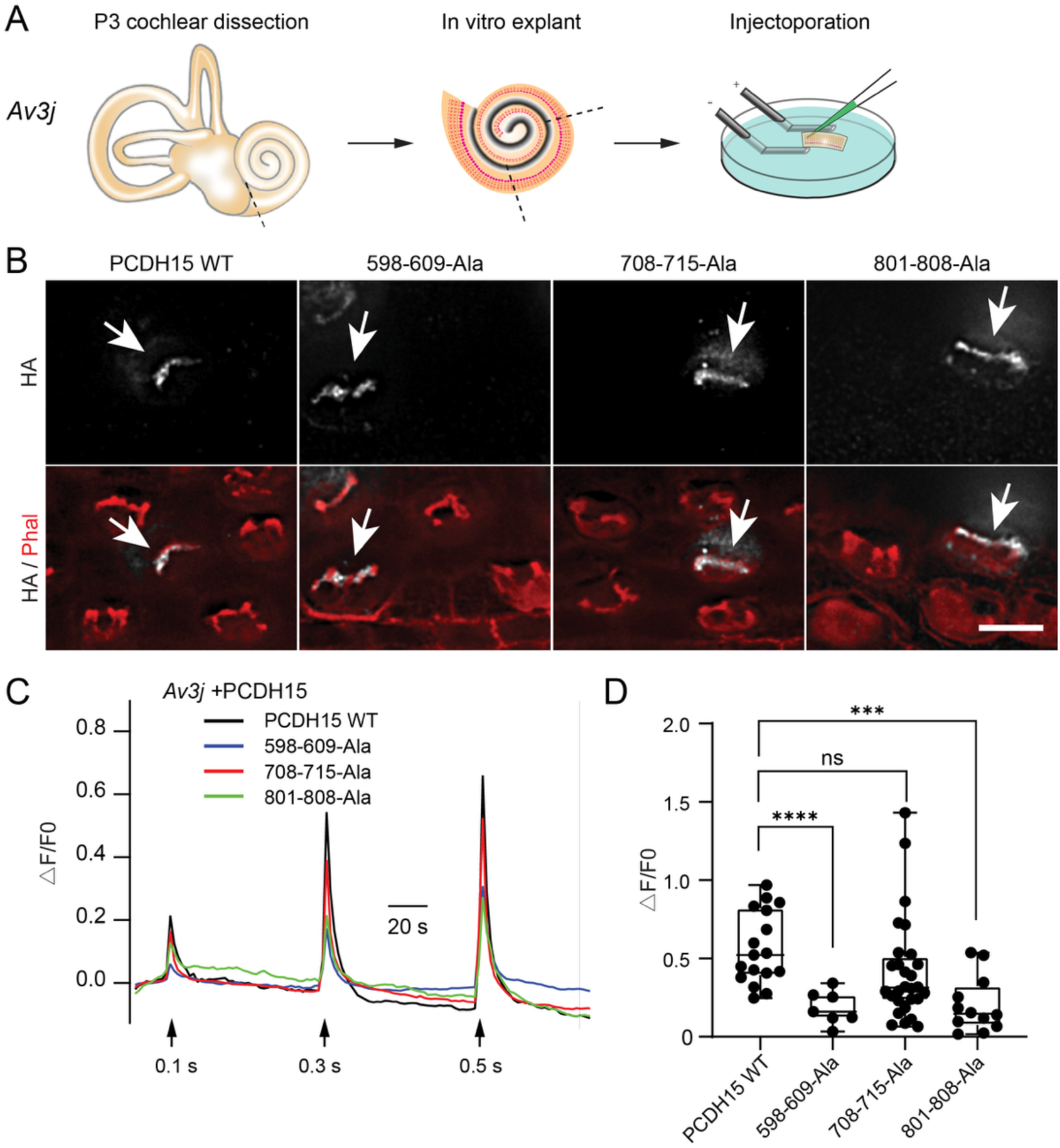
Rescue of MET in hair cells from PCDH15-deficient *av3J* mice. (A) Diagram of the experimental procedure: left, inner ear; middle, dissected cochlear sensory epithelium; right, cultured part of the cochlear sensory epithelium showing the injection needle and electroporation electrodes. (B) Immunofluorescence images of cochlear whole mounts injectoporated at P2-P3 with the indicated plasmids and stained after 1-2 day(s) in culture with phalloidin (red) to reveal stereocilia, and with antibodies to HA-tag (white) to detect PCDH15-HA. Arrows point to injectoporated cells expressing PCDH15 constructs. Scale bar: 5 μm. (C) Averaged GCaMP3 fluorescence signal traces. Intensity of fluid-jet stimulation was increased by changing pulse duration (from 0.1 sec, to 0.3 sec, to 0.5 sec). For quantitative analysis, the amplitude of the 2^nd^ Ca^2+^ response peak was measured. (D) Quantification of peak amplitude in OHCs from control and *av3J/av3J* mice. All data are mean ± SEM. One-way ANOVA Dunnett’s multiple comparisons test was used to determine statistical significance (***p < 0.001, ****p < 0.0001).

## Discussion

Here we provide new insights into the structure and function of PCDH15. Cryo-EM analysis shows that the first seven EC repeats of PCDH15 assemble into a right-handed double-helix homodimer. This architecture is in striking agreement with earlier electron micrographs of tip links in hair cells in vivo, which suggested that tip links consist of two strands that form a tightly interwound helix (24). Our structures provide a molecular view of PCDH15 that reveals multiple dimerization interfaces. In addition to the previously reported contact at the EC2-EC3 linker (10, 12), we identified lateral contacts at EC4-EC5 and EC6-EC7. The EC4-EC5 interface arises from a near-parallel alignment of the two protomers, whereas the strands cross over at the EC2-EC3 and EC5-EC6 linkers. Earlier work also identified a lateral interaction in the membrane-proximal region, where two PICA domains align (23). Together, these observations indicate that PCDH15 dimerization is mediated by several distinct interfaces, which likely cooperate to stabilize the helical dimer under the mechanical load experienced by tip links.

Previous data suggested that the membrane-proximal region containing the PICA domain can act as a hinge, permitting limited deflection from a straight conformation (23). These data also showed a split conformation of the two strands in some images. Notably, the PCDH15-EC8-PICA fragment lacked the crossover at EC6-EC7 (23), which our current structure indicates is likely important for lateral stabilization of the two strands within tip links and may restrain a split conformation.

Our functional data indicate that dimerization is important for PCDH15 function in MET. Although the cryo-EM resolution did not permit direct visualization of side-chain interactions, it was sufficient to identify interaction surfaces for targeted mutagenesis. Mutations in single dimerization interfaces had only modest effects on co-immunoprecipitation with full-length PCDH15, suggesting redundancy among the multiple contacts. In the isolated EC1-7 fragment, however, the same mutations noticeably weakened but did not abolish dimer formation, consistent with additive homodimer stabilization by multiple interfaces. When introduced into full-length PCDH15 and expressed in hair cells from PCDH15-deficient mice, these mutants trafficked to stereocilia but failed to fully restore MET, indicating that lateral interactions throughout the extracellular domain are critical for normal tip-link function, and that the TM and CTD domains also stabilize the PCDH15 homodimer.

Our results fit into the broader picture of tip-link mechanics. Tip links have been proposed to act as gating springs for the MET channel, and recent biophysical measurements using high-speed optical trapping on recombinant PCDH15 support this idea (22). Those experiments show that PCDH15’s mechanical behavior is Ca^2+-^dependent and that the molecule can undergo limited unfolding events at physiological Ca^2+^ concentrations, implying that its stiffness can be modulated. Crystallographic data indicate that linkers between EC3-EC4 and EC9-EC10 bind only two Ca2^+^ ions and that EC5-EC6 binds a single Ca^2+^ ion (9, 16, 18), which could locally increase flexibility. The helix like structure of the tip link also suggests that the dimer may act as a spring-like molecule. The two crossover points we observe may provide mechanical reinforcement under strain, while interposed regions with fewer Ca^2+^ ligands could permit controlled compliance. Computational studies similarly predict reduced rigidity in specific linkers (9). Whether PCDH15 adopts kinked conformations in vivo remains uncertain. Such conformations are observed in vitro (9) but may depend on the absence of mechanical load.

In summary, we describe a right-handed helical dimer architecture for PCDH15, identify two previously unrecognized lateral dimerization regions, and show that these lateral interactions are essential for PCDH15’s role in MET. Past studies have used molecular dynamics simulation to analyze the extent to which tip links might act as gating springs for MET channels. Our structural data provide additional constrains and can inform parameters for future simulation studies.

## Materials and Methods

### Plasmids construction for protein expression and Co-immunoprecipitation

Mus musculus (mm) PCDH15-CD2 full length (1-1781 amino acids) plasmid has been described in Webb et al., 2011. Mus musculus (mm) CDH23-EC1-3-FC (1-352 amino acids) plasmid has been described in Kazmierczak et al., 2007. Mus musculus (mm) DNA sequences encoding protein mmPCDH15 full-length, and fragments mmPCDH15 EC1-PICA (1-1381 amino acids), and mmPCDH15 EC1-7(1-824 amino acids) were subcloned into the pcDNA3.1 vector (ThermoFisher, V79520) with a Strep tag (WSHPQFEK), HA tag (YPYDVPDYA), or FLAG tag (DYKDDDDK) at the C-terminus, followed by a stop codon. Similarly, mmCDH23 EC1-3 (1-352 amino acids) was subcloned into pcDNA3.1, followed by an Fc tag and a Strep tag (WSHPQFEK) or a FLAG tag (DYKDDDDK) at the C-terminus, followed by a stop codon. PCDH15 truncated mutants and point mutants, including PCDH15-Δ598–609 mutant, PCDH15-Δ710–720 mutant, PCDH15-Δ801–808 mutant, PCDH15-598–609A, PCDH15-708-715A, PCDH15-801–808A, were introduced using the QuickChange II XL site-directed mutagenesis kit (Agilent). All constructs were validated by DNA sequencing.

Primers for the addition of a Strep-tag to CDH23-F3FC:

5’-TGGAGCCACCCGCAGTTCGAAAAATGACTCGAGCATGCATCTAGAG-3’

5’-TCCTGATTTACCCGGAGACAGGGA-3’

Primers for the addition of a FLAG-tag to CDH23-F3FC:

5’-GACTACAAGGACGACGATGACAAGTGACTCGAGCATGCATCTAGAGG-3’

5’-TCCTGATTTACCCGGAGACAGGGA-3’

Primers for the addition of a HA-tag to PCDH15-CD2:

5’-TACCCATACGATGTTCCAGATTACGCTTGATCGGCCGCGACTCTAGATC-3’

5’-AAGGGCTGTGTTGTAACCTTCGGA-3’

Primers for the addition of a FLAG-tag to PCDH15-CD2:

5’- GACTACAAGGACGACGATGACAAGTAACTAGAGGGCCCGTTTAAACCCG-3’

5’-AAGGGCTGTGTTGTAACCTTCGGA-3’

Primers for generating PCDH15-EC1-7-FLAG from PCDH15-CD2-FLAG: 5’-GACTACAAGGACGACGATGACAAG-3’

5’-AAAAACAGGACTGTTATCATCAATGTCCA-3’

Primers for generating PCDH15-EC1-7-HA from PCDH15-CD2-HA: 5’-TACCCATACGATGTTCCAGATTACG-3’

5’-AAAAACAGGACTGTTATCATCAATGTCCA-3’

Primers for introducing the 598–609 mutation in PCDH15-CD2:

5’-CAGCGGCCGCTGCAGCTGCAGCGGCCGCTGCAGCTGTGCTTCCTCCTAACAAC AACCAGAG-3’

5’-CTGCAGGCGGGGCGTTGT-3’

Primers for introducing the 708-715 mutation in PCDH15-CD2:

5’- GCCAACGCAGCGACGGACGCCAATGACAACGCTCCCGTGTTC-3’

5’-AGTGGCAGTTGAGGTTCCATC-3’

Primers for introducing the 801-808 mutation in PCDH15-CD2:

5’-GCGGCCGCTGCAGCTGCAGCGGCTACACTGTACATCAAGGTGTTGGACATTG-3’

5’-TGCTCCATCTGTTGCCACGAC-3’

Primers for deleting amino acids 598–609 in PCDH15-CD2: 5’-GTGCTTCCTCCTAACAACCAGAG-3’

5’-TGCAGGCGGGGCGTT-3’

Primers for deleting amino acids 710-720 in PCDH15-CD2: 5’-GTGTTCGATCCCTATCTGCCCAG-3’

5’-GTTCACAGTGGCAGTTGAGGTTC-3’

Primers for deleting amino acids 801-809 in PCDH15-CD2:

5’-CTGTACATCAAGGTGTTGGACATTGATG-3’

5’-TGCTCCATCTGTTGCCACGAC-3’

Primers for generating PCDH15-EC1-PICA-HA from PCDH15-CD2-HA:

5’- TCAGGAGACTACAAGGACGACGATGACAAGTAACTAGAGGGCCCGTTTAAACCCG-3’

5’-GAAGCGAGGAGAAAGCTTGGGG-3’

Primers for generating PCDH15-EC1-PICA-FLAG from PCDH15-CD2-FLAG:

5’- TCAGGAGACTACAAGGACGACGATGACAAGTAACTAGAGGGCCCGTTTAAAC CCG-3’

5’-GAAGCGAGGAGAAAGCTTGGGG-3’

### Protein expression and purification for gel filtration and cryo-EM

All PCDH15 and CDH23 fragments were expressed by transiently transfecting FreeStyle™ 293F cells (Thermo, R79007) or Expi293F GnTI- cells (Gibco, A39240) using Polyethylenimine “Max” (PEI MAX), MW 40,000 (Polysciences, 49553-93-7). Cells were harvested 48 hours post-transfection by centrifugation (3,000 g, 10 min, 4°C), washed with PBS (pH 7.4) containing protease inhibitors (0.8 µM aprotinin, 2 µg/mL leupeptin, 2 µM pepstatin A, and 1 mM phenylmethylsulfonyl fluoride), and pelleted again (3,800 g, 10 min, 4°C). Cells were lysed on ice using a blunt probe sonicator (3 cycles, 1 sec on, 1 sec off, for 1 min, 20 W power). The lysate was centrifuged to remove large cellular debris (3,800 g, 20 min, 4°C), and the solubilized protein was incubated overnight at 4°C with rotation using 0.75 mL Strep-Tactin® Sepharose® resin (2-1201-010) or Anti-DYKDDDDK G1 Affinity Resin (L00432) per 1 L of culture. The following day, the resin was collected via gravity flow, washed with 20 column volumes of lysis buffer (20 mM Tris, pH 8.0, 150 mM NaCl, 3 mM CaCl₂), and eluted using elution buffer containing either 10 mM D-desthiobiotin or 200 µg/mL DYKDDDDK synthetic peptide (Sino Biological, PP101274) in binding buffer. The eluate was collected in a centrifugal concentrator and concentrated to 500 µL at 4°C. If needed, the protein was further purified using a Superose 6 Increase 10/300 column (Cytiva, 29091596) on an ÄKTA FPLC. Peak fractions were collected and concentrated for subsequent experiments.

### Analytical size exclusion chromatography (SEC)

The Superose 6 Increase 10/300 GL column (Cytiva, 29091596) was pre-equilibrated with buffer (20 mM Tris, pH 8.0, 150 mM NaCl, 3 mM CaCl₂) on an FPLC system. Purified PCDH15 EC1-PICA, PCDH15 EC1-7, CDH23 EC1-3, CDH23 EC1-11, or their mixtures were analyzed using the same column equilibrated with the same buffer. All experiments were conducted at 4°C, and protein elution was monitored by absorbance at 280 nm (A280).

### Cryo-EM sample preparation and data collection

Graphene 300-mesh R 1.2/1.3 grids (Shuimu BioSciences) were plasma-treated using a Pelco Easiglow system (15 mA, 40 s glow time, 10 s hold time; Ted Pella, 91000). A 3 µL sample was applied to glow-discharged grids in an FEI Vitrobot Mark IV (Thermo Fisher Scientific) under the following conditions: 8°C, 100% humidity, 10 s wait time, blot force 0, and 3 s blot time. The grids were then plunge-frozen in liquid ethane. Imaging was performed using a 300-kV Titan Krios 3i microscope equipped with fringe-free imaging, a Falcon 4i camera, and a Selectris energy filter set to a 10 eV slit width. Micrographs were acquired at a dose rate of 7.11 e⁻ per pixel per second with a total dose of 40.00 e⁻ per Å². A total of 200 micrographs were collected for PCDH15 EC1-PICA, and 5,371 micrographs were collected for PCDH15 EC1-7. Movies were recorded using EPU software (Thermo Fisher Scientific) with a calibrated pixel size of 1.2 Å per pixel and a defocus range from -0.5 µm to -2.4 µm. Data collection was performed automatically using EPU software.

### Image processing

All aspects of image processing were performed using CryoSPARC(30). Refer to Figures S1-2 and Table S1 for details. Reconstruction quality was assessed for anisotropic contributions to the Fourier shell correlation (FSC) using 3DFSC (31).

### Model building, refinement, and structural analysis

All molecular modeling, refinement, and analysis were performed using UCSF ChimeraX (33), ChimeraX (34), ISOLDE (35), Coot (36, 37), and Phenix (38), accessed via the SBGrid consortium (39). The initial model of PCDH15 EC1-7 was predicted using AlphaFold3^14^. Each cadherin domain was extracted from the predicted model and rigid-body fitted into the composite map. ISOLDE was then used to refine individual amino acid positions relative to their respective locally refined maps (35). Further real-space refinement and geometry minimization were performed using Phenix, while amino acid and sidechain rotamer positions were manually adjusted in (36, 37). MolProbity (40) was used for model validation, and final model visualizations were generated using ChimeraX (34).

### Co-immunoprecipitation and western blotting

Co-immunoprecipitations (Co-IPs) were performed as previously described (41–43). Briefly, HEK293 cells were transfected with plasmids using Lipofectamine 3000 (Thermo Fisher). After 36 hours, cells were lysed in a modified RIPA buffer containing 50 mM Tris (pH 8.0), 150 mM NaCl, 1% NP-40, 0.5% sodium deoxycholate, 0.1% sodium dodecyl sulfate (SDS), and a Roche Complete Protease Inhibitor Tablet. Lysates were incubated on a rotator at 4°C for 30 minutes, followed by centrifugation at 16,000 rpm for 20 minutes at 4°C. At this stage, 5% of the lysate was set aside as an input control, while the remainder was immunoprecipitated for 1 hour at 4°C using either EZ View Red anti-HA Affinity Gel (Sigma-Aldrich, Cat# E6779-1ML) or EZ View Red anti-FLAG Affinity Gel (Sigma-Aldrich, Cat# F2426-1ML). After immunoprecipitation, the affinity gel was washed three times with IP buffer (50 mM Tris, pH 8.0, 150 mM NaCl, 1% NP-40). The remaining buffer was discarded, and proteins were eluted using 1X SDS Sample Buffer (Bio-Rad) containing β-mercaptoethanol (BME, Bio-Rad). Eluted immunoprecipitated proteins and input lysates were separated on Novex™ WedgeWell™ 4–20% Tris-Glycine Mini Protein Gels (Life Technologies) and transferred onto PVDF membranes (Sigma) at 250 mA for 2.5 hours at 4°C using the Mini Blot Module (Bio-Rad) with Bio-Rad transfer buffer containing 20% methanol. Membranes were blocked for 1 hour in 5% Blotting-Grade Blocker Non-Fat Dry Milk (Bio-Rad) in 1X TBST (20 mM Tris-HCl, pH 7.5, 150 mM NaCl, 0.05% Tween-20), followed by overnight incubation at 4°C with primary antibodies (see below) in 5% milk in 1X TBST. The next day, membranes were washed three times in 1X TBST, incubated for 1 hour at room temperature with HRP-conjugated secondary antibodies in 5% milk in 1X TBST, and washed three more times with 1X TBST. Signals were detected using Clarity ECL Substrate (Bio-Rad) on a G-Box ECL imager (Syngene).

Co-IP quantification was performed using ImageJ and GraphPad Prism 6. Band intensities of IP and Co-IP proteins were measured using ImageJ for both immunoprecipitated and whole-cell lysate proteins. Co-IP intensity values were normalized to account for expression levels and immunoprecipitation efficiency, then divided by controls to generate relative Co-IP intensity values. Mean relative Co-IP intensity values were calculated by averaging across independent experiments, and statistical significance was assessed using Student’s t-test.

Primary Antibodies: Rabbit anti-FLAG (1:500, Cell Signaling, Cat# 14793S), Rabbit anti-FLAG (1:500, Cell Signaling, Cat# F7425), Rabbit anti-HA (1:500, Cell Signaling, Cat# 3724). Secondary Antibodies: HRP-linked anti-rabbit IgG

### Injectoporation of cochlear hair cells

Injectoporation experiments were performed to express exogenous DNA in cochlear hair cells, as previously described (12, 27–29). Briefly, the organ of Corti was isolated from *Pcdh15*-deficient Ames waltzer (av3J) mice²³ and cultured in DMEM/F12 supplemented with 1.5 µg/mL ampicillin. Briefly, the organ of Corti was isolated from P2-3 *Pcdh15*-deficient Ames waltzer (av3J) mice²³ and cultured in DMEM/F12 supplemented with 1.5 µg/mL ampicillin. PCDH15-CD2 WT and point mutation plasmid DNA constructs (1 µg/µL) together with GCamp3 construct (0.5 µg/µL) were injected into explants using glass pipettes (2 µm diameter) positioned between rows of outer hair cells. Explants were then electroporated with four pulses at 60 V (15 msec pulse length, 1 sec inter-pulse intervals) using an ECM 830 square wave electroporator (BTX). Following electroporation, the culture medium was replaced with DMEM/F12 containing 10% fetal bovine serum (FBS) and 1.5 µg/mL ampicillin. Explants were incubated for 1–2 days at 37°C with 5% CO₂ before calcium imaging and/or immunostaining.

### Whole mount Immunostaining and Imaging and PCDH15 expression level quantification

Whole-mount cochlear explant preparations were fixed and immunostained as previously described (41–43). Briefly, culture dishes containing explants were incubated in 1× HBSS with 4% PFA and 0.1 mM CaCl₂ for 30 minutes at 4°C, followed by three washes in PBS. The Organ of Corti was permeabilized in PBS containing 0.5% Triton X-100 for 30 minutes at 4°C with gentle agitation. After permeabilization, tissues were blocked in PBS containing 10% goat serum (GS) for 4–6 hours at room temperature (RT). Samples were then incubated overnight at 4°C in PBS with 5% GS and primary antibodies (Rabbit anti-HA, 1:200; cell signaling, RRID: AB_1549585) Following primary antibody incubation, tissues were washed three times in PBS and incubated for 1 hour at RT in PBS with 5% GS, secondary antibodies (Goat anti-Rabbit IgGF(ab’)2, AlexaFluor 555, 1:5000; Invitrogen Cat#:A-21430;RRID:AB_2535851), and fluorescently conjugated phalloidin (Life Technologies, Phalloidin 405, 1:1000) to label stereocilia. After secondary antibody incubation, samples were washed three times in PBS, mounted using ProLong Gold (Invitrogen), and imaged using a wide-field fluorescence deconvolution microscope with 100× objectives (Deltavision, GE Life Sciences).

### Calcium imaging

Calcium imaging was performed as previously described (27, 28) using G-CaMP3 as a Ca²⁺ indicator. Imaging was conducted on an upright Olympus BX51WI microscope equipped with a 60× water-immersion objective and a Qimaging ROLERA-QX camera, controlled by Micro-Manager 1.3 software. Hair bundles were stimulated using a fluid jet applied through a glass electrode (2 µm tip diameter) filled with bath solution. Stimuli were delivered using Patchmaster 2.35 software (HEKA) and a 20 psi air pressure pulse generated by a Picospritzer III microinjector. Images were collected at a 2-second sampling rate, and fluid-jet stimulations were applied at 0.1, 0.3, and 0.5 seconds with 60-second intervals between each stimulation. Responses induced by 0.3-second stimulations were used for quantitative analysis. Data analysis was performed using Excel (Microsoft) and Igor pro 6 (WaveMetrics, Lake Oswego, OR). Calcium signal (ΔF/F) was calculated with the equation: (F-F0)/F0, where F0 is the averaged fluorescence baseline at the beginning. All data are mean ± SEM. Student’s two-tailed unpaired t test was used to determine statistical significance (*p < 0.05, **p < 0.01, ***p < 0.001)

Plasmids used for injectoporation were as follows: (i) CMV-GCaMP3 has been described previously(27, 28); (ii) CMV-PCDH15-CD2 (26) was used to express wild-type PCDH15 in hair cells. All PCDH15 point mutations were generated using the Quick-Change II XL site-directed mutagenesis Kit from Agilent.

## Data availability

The accession code for PCDH15-EC1-7 is EMDB: EMD-75730. The primary cryo-EM maps (before local refinement and signal subtraction) are included in each deposition, while the composite map and locally refined maps are provided as supplemental files. The PCDH15-EC1-7 structure is available in the Protein Data Bank (PDB) under accession code PDB: 11IY. Source data are provided with this paper.

## Acknowledgments

The authors like to thank members of the Müller and Twomey laboratories for discussions and input to the project. This work was funded by the NIH (RO1DC005965 to UM; R35GM154904 to E.C.T; F31GM157915 to L.D.); by the Fondation Pour l’Audition (FPA RD-2024-1 to UM); by the David M. Rubenstein Fund for Hearing Research (to U.M.); by the Searle Scholars Award via the Kinship Foundation (to E.C.T.). U.M. is a Bloomberg Distinguished Professor for Neuroscience and Biology. All cryo-EM work was performed at the Beckman Center for Cryo-EM at Johns Hopkins.

---

**Figure S1.**
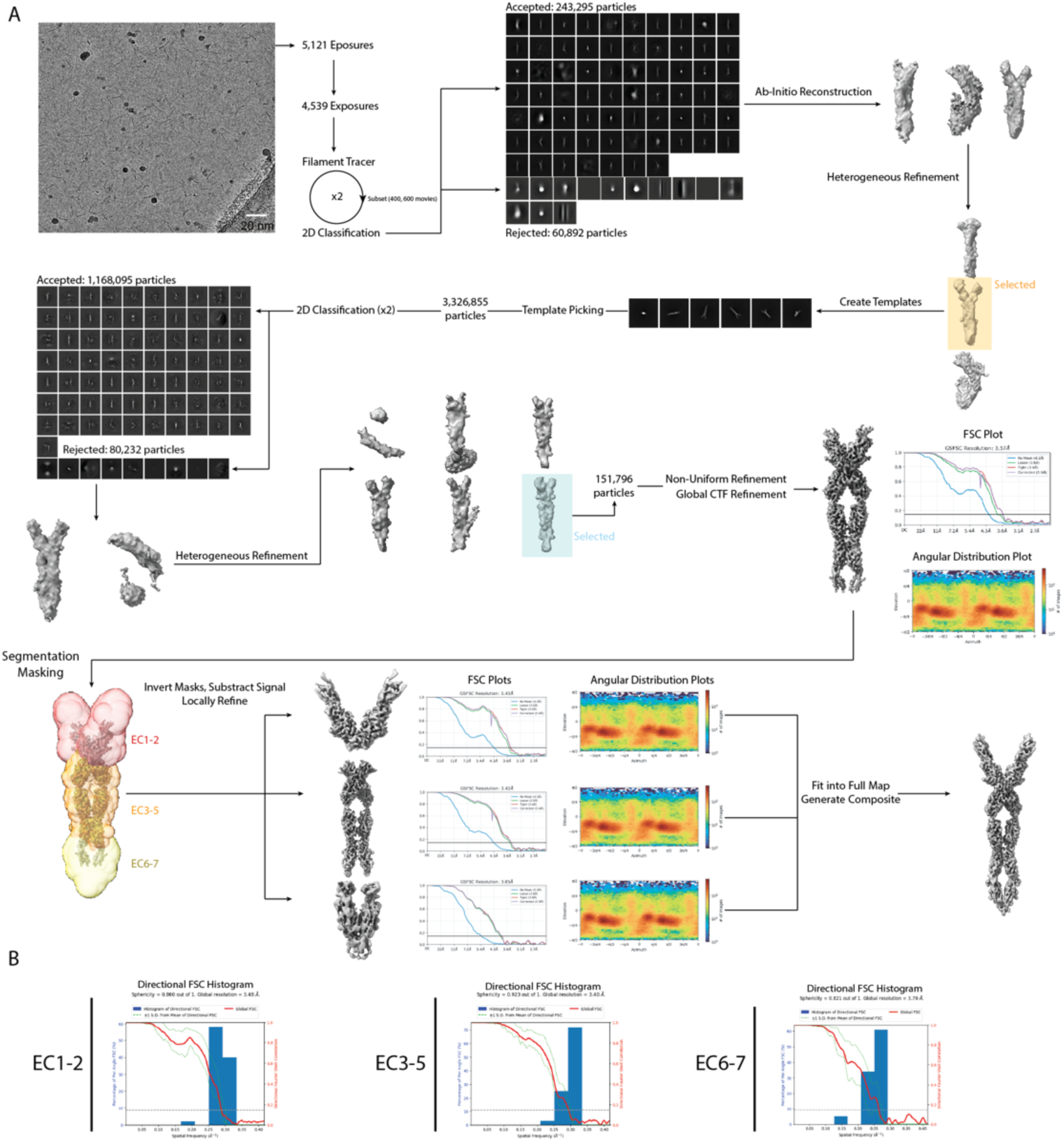
Image processing workflow for Pcdh15 EC1-7. (A) Processing details from cryoSPARC. Each final map includes the Fourier shell correlation (FSC) plots, with resolution calculated at FSC=0.143 and an angular distribution plot for particle images contributing to the reconstruction. (B) Three-dimensional FSC (3DFSC) calculated for each local map in panel A.

**Figure S2.**
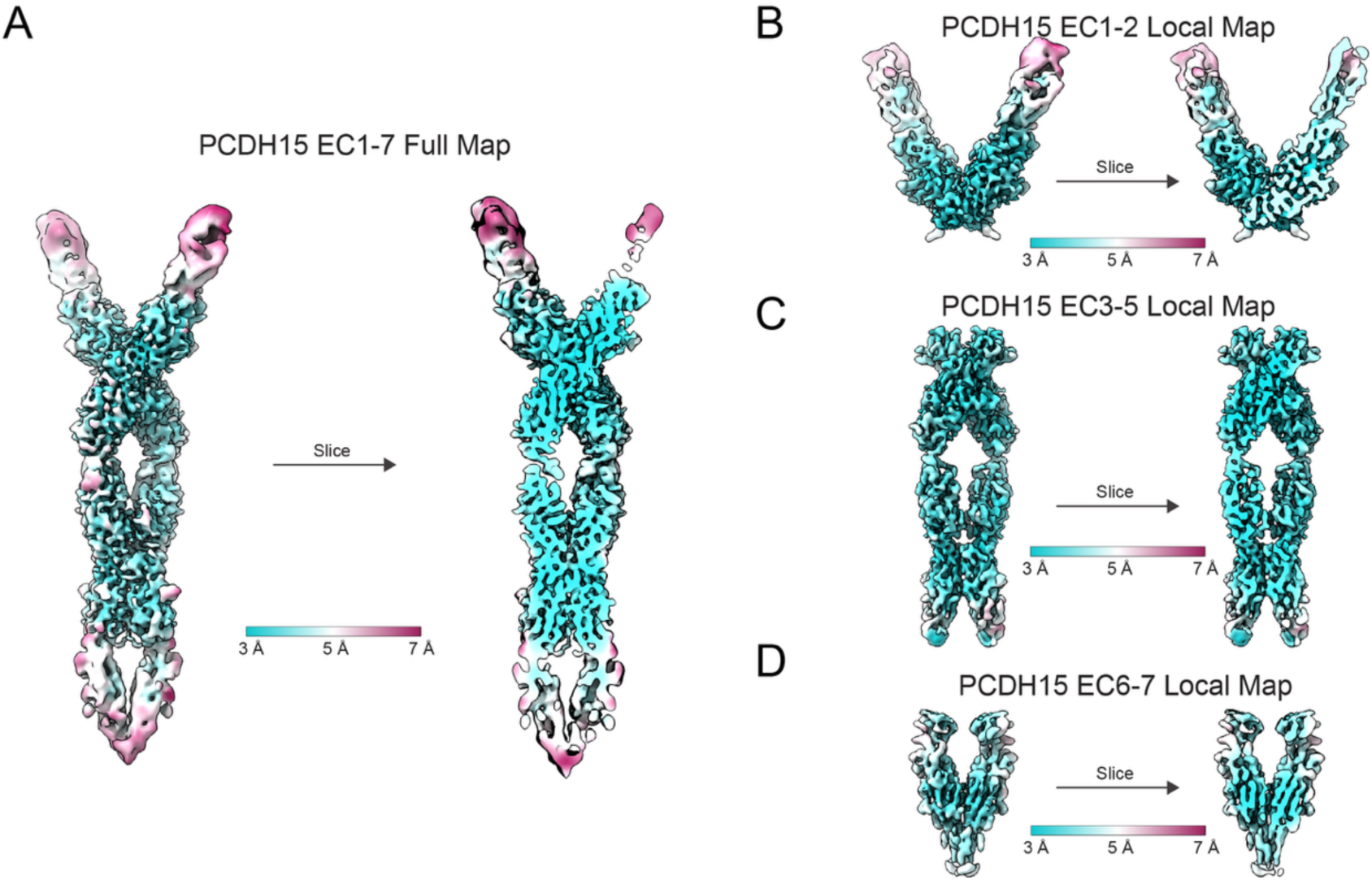
Local resolution of cryo-EM maps. The maps are colored according to local resolution for the full EC1-7 map (A), EC1-2 (B), EC3-5 (C), and EC6-7 (D). Resolution was calculated at each voxel according to FSC=0.143. For each map, the whole map is shown (left) and a central slice (right).

**Table S1.**
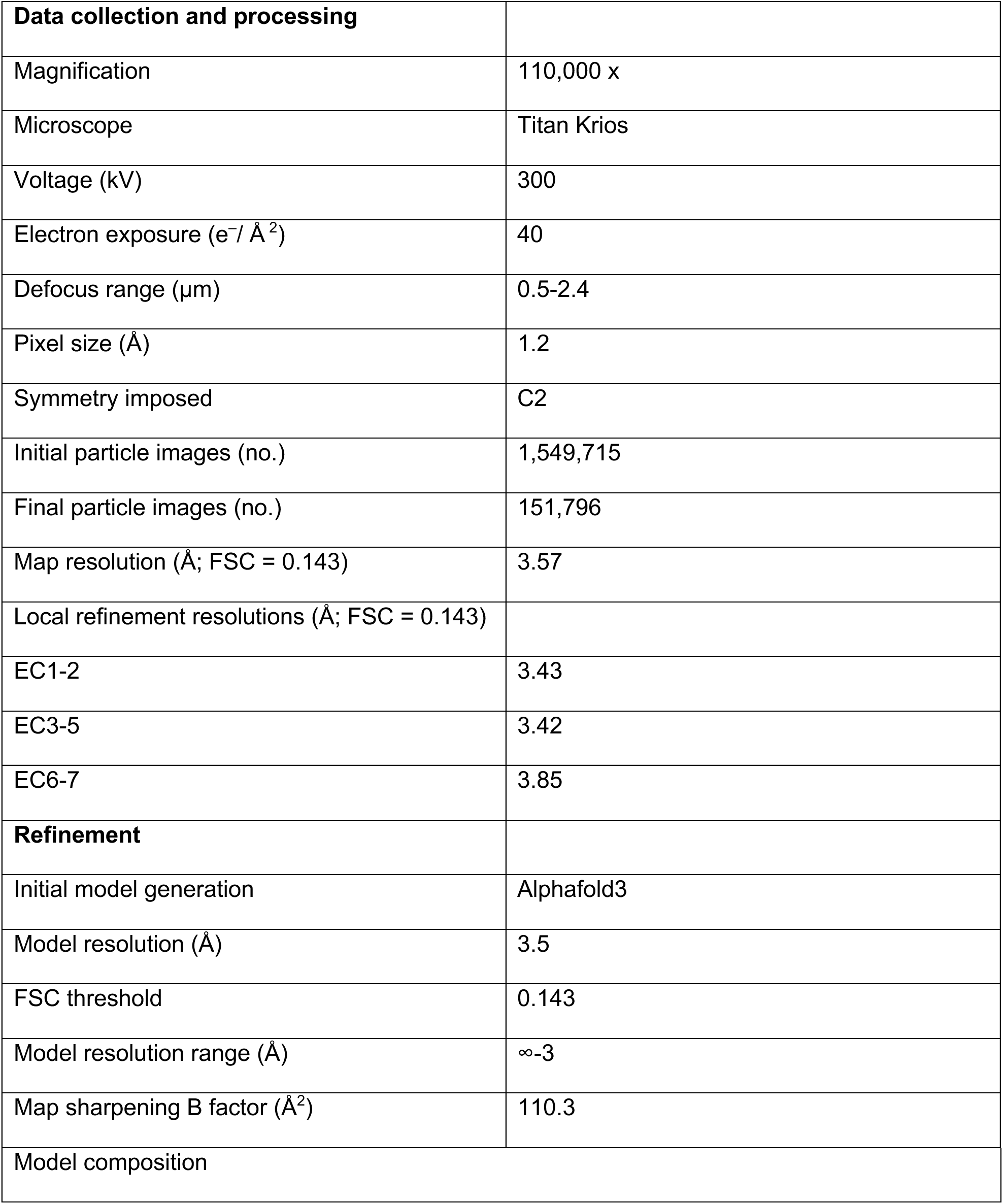
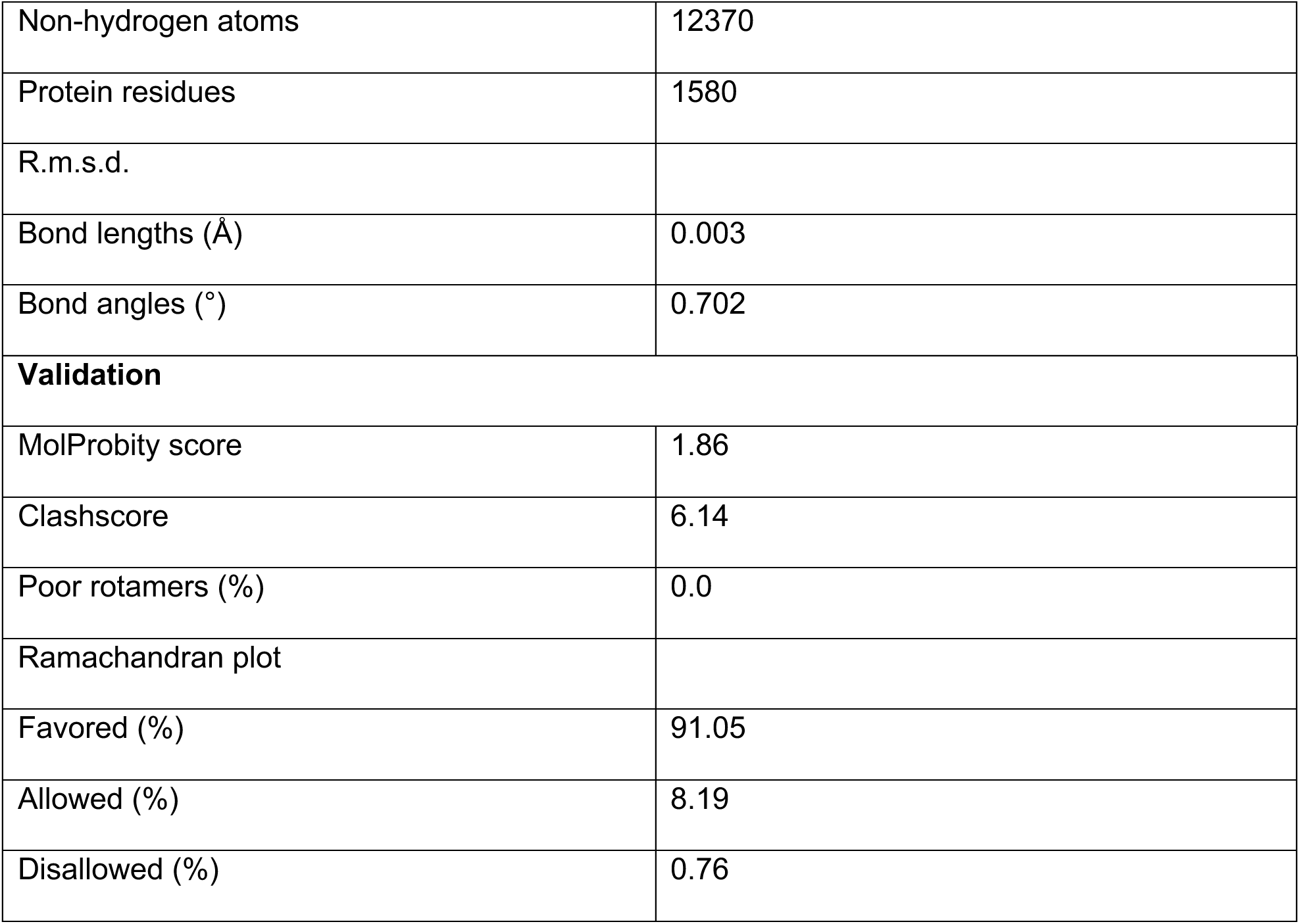
Cryo-EM data collection, refinement, and validation statistics.

## References

1. R. Fettiplace, Hair Cell Transduction, Tuning, and Synaptic Transmission in the Mammalian Cochlea. Compr Physiol 7, 1197–1227 (2017).

2. X. Qiu, U. Muller, Sensing sound: Cellular specializations and molecular force sensors. Neuron 110, 3667–3687 (2022).

3. P. Kazmierczak et al., Cadherin 23 and protocadherin 15 interact to form tip-link filaments in sensory hair cells. Nature 449, 87–91 (2007).

4. Z. M. Ahmed et al., Mutations of the protocadherin gene PCDH15 cause Usher syndrome type 1F. Am J Hum Genet 69, 25–34 (2001).

5. K. N. Alagramam et al., Mutations in the novel protocadherin PCDH15 cause Usher syndrome type 1F. Hum Mol Genet 10, 1709–1718 (2001).

6. H. Bolz et al., Mutation of CDH23, encoding a new member of the cadherin gene family, causes Usher syndrome type 1D. Nat Genet 27, 108–112 (2001).

7. J. M. Bork et al., Usher syndrome 1D and nonsyndromic autosomal recessive deafness DFNB12 are caused by allelic mutations of the novel cadherin-like gene CDH23. Am J Hum Genet 68, 26–37 (2001).

8. F. Di Palma et al., Mutations in Cdh23, encoding a new type of cadherin, cause stereocilia disorganization in waltzer, the mouse model for Usher syndrome type 1D. Nat Genet 27, 103–107 (2001).

9. R. Araya-Secchi, B. L. Neel, M. Sotomayor, An elastic element in the protocadherin-15 tip link of the inner ear. Nat Commun 7, 13458 (2016).

10. D. Choudhary et al., Structural determinants of protocadherin-15 mechanics and function in hearing and balance perception. Proc Natl Acad Sci U S A 117, 24837–24848 (2020).

11. P. De-la-Torre, D. Choudhary, R. Araya-Secchi, Y. Narui, M. Sotomayor, A Mechanically Weak Extracellular Membrane-Adjacent Domain Induces Dimerization of Protocadherin-15. Biophys J 115, 2368–2385 (2018).

12. G. Dionne et al., Mechanotransduction by PCDH15 Relies on a Novel cis-Dimeric Architecture. Neuron 99, 480–492 e485 (2018).

13. H. M. Elledge et al., Structure of the N terminus of cadherin 23 reveals a new adhesion mechanism for a subset of cadherin superfamily members. Proc Natl Acad Sci U S A 107, 10708–10712 (2010).

14. A. Jaiganesh et al., Zooming in on Cadherin-23: Structural Diversity and Potential Mechanisms of Inherited Deafness. Structure 26, 1210–1225 e1214 (2018).

15. Y. Narui, M. Sotomayor, Tuning Inner-Ear Tip-Link Affinity Through Alternatively Spliced Variants of Protocadherin-15. Biochemistry 57, 1702–1710 (2018).

16. R. E. Powers, R. Gaudet, M. Sotomayor, A Partial Calcium-Free Linker Confers Flexibility to Inner-Ear Protocadherin-15. Structure 25, 482–495 (2017).

17. M. Sotomayor, W. A. Weihofen, R. Gaudet, D. P. Corey, Structural determinants of cadherin-23 function in hearing and deafness. Neuron 66, 85–100 (2010).

18. M. Sotomayor, W. A. Weihofen, R. Gaudet, D. P. Corey, Structure of a force-conveying cadherin bond essential for inner-ear mechanotransduction. Nature 492, 128–132 (2012).

19. T. J. Boggon et al., C-cadherin ectodomain structure and implications for cell adhesion mechanisms. Science 296, 1308–1313 (2002).

20. O. J. Harrison et al., Two-step adhesive binding by classical cadherins. Nat Struct Mol Biol 17, 348–357 (2010).

21. S. D. Patel et al., Type II cadherin ectodomain structures: implications for classical cadherin specificity. Cell 124, 1255–1268 (2006).

22. T. F. Bartsch et al., Elasticity of individual protocadherin 15 molecules implicates tip links as the gating springs for hearing. Proc Natl Acad Sci U S A 116, 11048–11056 (2019).

23. J. Ge et al., Structure of mouse protocadherin 15 of the stereocilia tip link in complex with LHFPL5. Elife 7 (2018).

24. B. Kachar, M. Parakkal, M. Kurc, Y. Zhao, P. G. Gillespie, High-resolution structure of hair-cell tip links. Proc Natl Acad Sci U S A 97, 13336–13341 (2000).

25. E. Pepermans et al., The CD2 isoform of protocadherin-15 is an essential component of the tip-link complex in mature auditory hair cells. EMBO Mol Med 6, 984–992 (2014).

26. S. W. Webb et al., Regulation of PCDH15 function in mechanosensory hair cells by alternative splicing of the cytoplasmic domain. Development 138, 1607–1617 (2011).

27. W. Xiong et al., TMHS is an integral component of the mechanotransduction machinery of cochlear hair cells. Cell 151, 1283–1295 (2012).

28. W. Xiong, T. Wagner, L. Yan, N. Grillet, U. Muller, Using injectoporation to deliver genes to mechanosensory hair cells. Nat Protoc 9, 2438–2449 (2014).

29. B. Zhao et al., TMIE is an essential component of the mechanotransduction machinery of cochlear hair cells. Neuron 84, 954–967 (2014).

30. A. Punjani, J. L. Rubinstein, D. J. Fleet, M. A. Brubaker, cryoSPARC: algorithms for rapid unsupervised cryo-EM structure determination. Nat Methods 14, 290–296 (2017).

31. Y. Z. Tan et al., Addressing preferred specimen orientation in single-particle cryo-EM through tilting. Nat Methods 14, 793–796 (2017).

32. B. A. Barad et al., EMRinger: side chain-directed model and map validation for 3D cryo-electron microscopy. Nat Methods 12, 943–946 (2015).

33. E. F. Pettersen et al., UCSF Chimera--a visualization system for exploratory research and analysis. J Comput Chem 25, 1605–1612 (2004).

34. E. F. Pettersen et al., UCSF ChimeraX: Structure visualization for researchers, educators, and developers. Protein Sci 30, 70–82 (2021).

35. T. I. Croll, ISOLDE: a physically realistic environment for model building into low-resolution electron-density maps. Acta Crystallogr D Struct Biol 74, 519–530 (2018).

36. P. Emsley, K. Cowtan, Coot: model-building tools for molecular graphics. Acta Crystallogr D Biol Crystallogr 60, 2126–2132 (2004).

37. P. Emsley, B. Lohkamp, W. G. Scott, K. Cowtan, Features and development of Coot. Acta Crystallogr D Biol Crystallogr 66, 486–501 (2010).

38. D. Liebschner et al., Macromolecular structure determination using X-rays, neutrons and electrons: recent developments in Phenix. Acta Crystallogr D Struct Biol 75, 861–877 (2019).

39. A. Morin et al., Collaboration gets the most out of software. Elife 2, e01456 (2013).

40. C. J. Williams et al., MolProbity: More and better reference data for improved all-atom structure validation. Protein Sci 27, 293–315 (2018).

41. C. L. Cunningham et al., TMIE Defines Pore and Gating Properties of the Mechanotransduction Channel of Mammalian Cochlear Hair Cells. Neuron 107, 126–143 e128 (2020).

42. X. Liang et al., CIB2 and CIB3 are auxiliary subunits of the mechanotransduction channel of hair cells. Neuron 109, 2131–2149 e2115 (2021).

43. X. Qiu, X. Liang, J. P. Llongueras, C. Cunningham, U. Muller, The tetraspan LHFPL5 is critical to establish maximal force sensitivity of the mechanotransduction channel of cochlear hair cells. Cell Rep 42, 112245 (2023).

